# Interplay of *Porphyromonas gingivalis* and *Aggregatibacter actinomycetemcomitans* along with Circulating miR-21 and miR-155 as Potential Biomarkers for Pancreatic Cancer

**DOI:** 10.1101/2025.05.21.655259

**Authors:** Zeinab Hesami, Meysam Olfatifar, Amir Sadeghi, Samira Mohammadi-Yeganeh, Hesameddin Eghlimi, Valerio Pazienza, Mojdeh Hakemi-Vala, Hamidreza Houri

## Abstract

The associations between oral bacterial pathogens and the risk of pancreatic cancer (PC) have been reported in several epidemiological studies. In this study, we evaluated the diagnostic potential of periodontal pathogens *Porphyromonas gingivalis* and *Aggregatibacter actinomycetemcomitans* in combination with circulating oncomiRNAs, including miR-21 and miR-155. A total of 41 PC patients and 40 age- and sex-matched controls were recruited for the study. The salivary bacterial load of *P. gingivalis* and *A. actinomycetemcomitans*, along with the copy number of miR-21 and miR-155 in blood, were measured using quantitative real-time PCR. Subsequently, the predictive power of the selected biomarkers in PC diagnosis was determined. Elevated load of the periodontal pathogens *P. gingivalis* in females (OR=2.31; 95% CI 0.98-5.47) and *A. actinomycetemcomitans* in diabetic individuals (OR=3.66; 95% CI 0.47-6.68) were associated with a higher risk of PC. Moreover, the diagnostic model incorporating two salivary species and two circulating miRNAs demonstrated an AUC of 0.878 (95% CI 0.802-0.955). This study offers compelling new evidence supporting the idea that the combined analysis of salivary microbiota and circulating miRNAs serves as an informative avenue for the discovery of non-invasive biomarkers for PC.

## Introduction

Pancreatic cancer (PC) is an extremely lethal disease, characterized by high mortality within five years of diagnosis. GLOBOCAN estimates projected that by 2040, the global incidence and mortality rates of PC could increase by over 75% and 80%, respectively (1, 2). The unfavorable prognosis of PC is largely due to its tendency to quickly spread to the lymphatic system and distant organs (3-5). The aggressive biological behavior of PC, coupled with its high resistance to conventional chemotherapies and the absence of effective sensitive diagnostic biomarkers, results in a 5-year survival rate of only 5% for individuals diagnosed with the disease (6, 7). Approximately 15% to 20% of PC patients present with a disease that is eligible for surgical resection. However, of these patients, only about 20% achieve a 5-year survival rate (4). PC stems from a complex interplay of etiological factors, with cigarette smoking being widely recognized as the only established modifiable risk factor. However, evidence also points to a potential link between diabetes, obesity, and insulin resistance with an elevated risk of developing this malignancy (8, 9). Furthermore, the strong correlation between chronic pancreatitis (CP) and a significantly heightened risk of PC suggests that inflammation may play a pivotal role in the initiation and progression of the disease. Inflammatory pathways may drive cellular proliferation and mutagenesis, hinder the ability to adapt to oxidative stress, promote angiogenesis, inhibit apoptosis, and elevate the secretion of pro-inflammatory signaling molecules (10).

The oral cavity serves as a significant reservoir of microorganisms, encompassing over 700 species, collectively known as the oral microbiome (11). Mounting evidence indicates that oral microbiota plays crucial roles in human health, influencing immune regulation, carcinogen metabolism, and nutrient processing (12). Periodontitis, an inflammatory condition of the oral cavity caused by dysbiosis of the oral microbiota, has been linked to an elevated risk of PC in several longitudinal studies (13-15). Additionally, recent epidemiological studies have shown that poor oral health and the presence of circulating and salivary antibodies to specific oral pathogens are associated with an increased risk of PC (13, 16, 17). Further prospective investigations have also underscored the potential of periodontal pathogens as biomarkers for the early detection of PC (18, 19).

MicroRNAs are short RNA molecules, typically 18-24 nucleotides in length, that have been evolutionarily conserved (20). These molecules regulate the stability and translation of target mRNAs by binding to complementary sequences in the 3’ untranslated regions (3’ UTRs). This regulatory mechanism is crucial for maintaining cellular homeostasis and supporting developmental processes. In the recent years, the significant role of miRNAs in controlling cell growth and their involvement in cancer has become increasingly apparent (21). Recently, numerous miRNAs have been identified in cell-free environments within various biofluids, including whole blood, plasma, serum, saliva, tears, urine, stool, pancreatic juice, breast milk, cerebrospinal fluid, and peritoneal and pleural fluids (22). These circulating miRNAs display specific expression patterns that correlate with changes in physiological conditions and the presence of disease. Numerous studies have identified various miRNAs with dysregulated expression linked to PC, highlighting their potential as circulating biomarkers for diagnosis of the disease at early stages (23). In this study, we conducted a comprehensive analysis comparing the load of *Porphyromonas gingivalis* and *Aggregatibacter actinomycetemcomitans* in salivary samples with the copy number of miR-21 and miR-155 in blood samples from both patients diagnosed with PC and control subjects, using quantitative real-time PCR. Additionally, we assessed the performance and translational potential of these periodontal pathogens and circulating miRNAs as potential biomarkers for the non-invasive detection of PC.

## Materials and Methods

### Study population and sample collection

This study was reviewed and approved by the Institutional Ethical Review Committee of the School of Medicine at Shahid Beheshti University of Medical Sciences (No.IR.SBMU.MSP.REC.1402.184). In this case-control study conducted between 2022 and 2024 at Taleghani Hospital in Tehran, Iran, participants were prospectively enrolled following standardized protocols for the collection, processing, and storage of biological samples. Individuals over 18 years of age with suspected PC, confirmed through pathology diagnosis during endoscopic ultrasound (EUS) procedures, were included as cases. Controls, matched for age, sex, and hospital, were chosen from orthopedic ward inpatients without hepatobiliary conditions or a history of cancer. Written informed consent was obtained from participating wards and study participants, respectively. Demographic data, including age, sex, body mass index (BMI), tobacco and opium use, etc., were collected during face-to-face interviews through a structured questionnaire. Clinical and paraclinical information was retrieved from hospital charts, including the stage of the disease, past medical history, past surgery history, social history, and laboratory findings. Before the sampling process, all the volunteers were instructed to not eat for 1 h prior to saliva sample collection. Unstimulated saliva samples were collected, aliquoted, and preserved in RNALater and stored at 4°C overnight, then stored at −70°C until DNA extraction. Blood samples were processed immediately on the same day of sample collection for RNA extraction and cDNA synthesis.

### Sample processing

Salivary samples were thawed, and DNA was extracted and purified using the E.Z.N.A.® MicroElute® Genomic DNA Kit (Omega Bio-tek, Georgia, USA). miRNA extraction from whole blood was performed using the SanPrep Column microRNA Miniprep Kit (Bio Basic, Markham, ON, Canada), followed by cDNA synthesis via RevertAid® RT enzyme (Invitrogen, Thermo Fisher Scientific) and RT stem-loop primers according to established protocols (24). The yield and purity of the isolated DNA and RNA were assessed using a spectrophotometer (Nanodrop, Thermo Fisher Scientific, Waltham, MA, USA).

### Bacterial strains

The standard strains of *P. gingivalis* (ATCC33277) and *A. actinomycetemcomitans* (ATCC24523) were obtained from the School of Public Health, Tehran University of Medical Sciences and Dental School, Shahid Beheshti of Medical Sciences, respectively. These strains were cultured on brucella agar supplemented with 10% sheep blood, hemin, and menadione under anaerobic conditions (5% CO2, 10% H2, 85% N2) at 37°C. Bacterial DNA extraction was carried out from purified colonies of each strain using the DNA QIAamp DNA Mini Kit (QIAGEN, Hilden, Germany) following the provided instructions.

### Serial dilution preparation and quantitative real-time PCR

For absolute quantification of the bacterial biomarker candidates, serial dilutions were prepared to obtain standard curves based on the genomic size of standard strains and the optical density (OD) of extracted DNA. In addition, synthesized nucleotide sequences of miR-21 and miR-155 were provided to prepare serial dilutions based on fragment size and molar mass per base pair.

Sequences of the oligonucleotide primers used for real-time qPCR are shown in Table S1. Real-time PCR was carried out in duplicate in reaction volumes of 10 µl using RealQ plus 2X Master Mix Probe High ROX (Ampliqon, Herlev, Denmark). The thermal cycling conditions consisted of an initial denaturing for 15 min at 95°C, followed by 40 cycles of amplification with three steps: 95°C for 20 seconds, then either 62°C for 30 seconds, and 72°C for 50 seconds for *P. gingivalis*, or 56°C for 30 seconds and 72°C for 40 seconds for *A. actinomycetemcomitans*. Additionally, miRNAs were amplified in two steps of 95°C for 15 s and 60°C for 50 s. All qPCR tests were performed using LightCycler® (Roche Life Science, Berlin, Germany).

### Statistical analysis

The significance of the bacterial load and copy number of the miRNAs was determined between the PC group and the non-cancer control group through an independent *t*-test. Association of two microbial data as well as copy number of two circulating miRNAs to subject status were assessed using the ‘adonis2’ implementation of PERMANOVA and distance-based redundancy analysis. To investigate the relationship between these biomarkers and PC risk, a least absolute shrinkage and selection operator (LASSO) logistic regression method was applied to estimate odds ratios (ORs) and their corresponding 95% confidence intervals (CIs). Features selected during the LASSO process were included in risk assessment analyses within stratified subgroups of cases and controls. Subsequently, receiver operating characteristic (ROC) curves were generated, and the area under the curve (AUC) was determined through numerical integration via both univariable and multivariable logistic regression models. All statistical analyses were conducted using R version 4.3.2.

## Results

### Characteristics of the Study Participants

As shown in Table 1, of the included PC patients (n=41), 68% were male and the majority of them were more likely to be current smokers (39%), and have type 2 diabetes (61%). Among patients diagnosed with PC, most of the tumors were well differentiated (G_1_) and located in the head of the pancreas (77.1%).

**Table 1:**
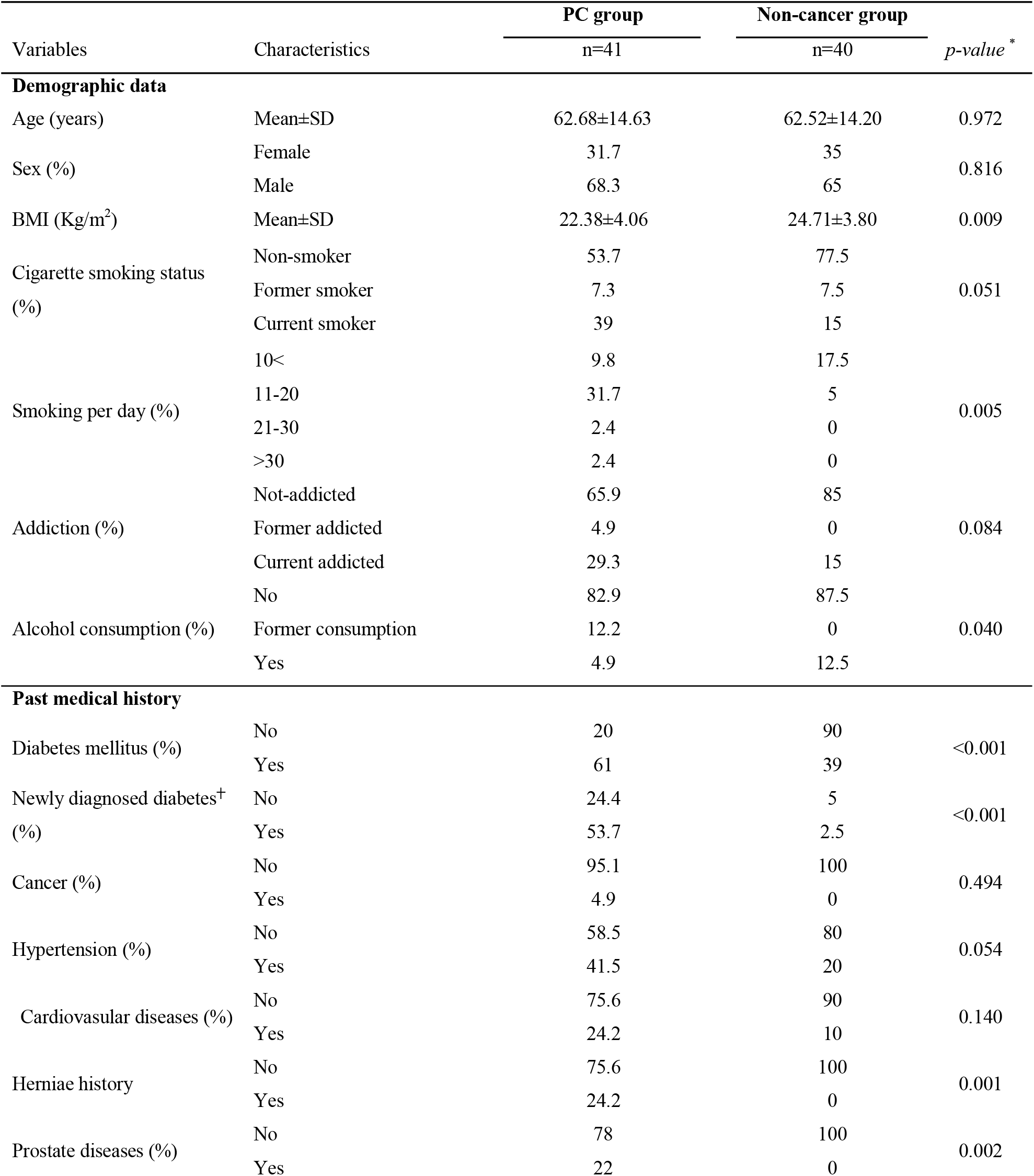
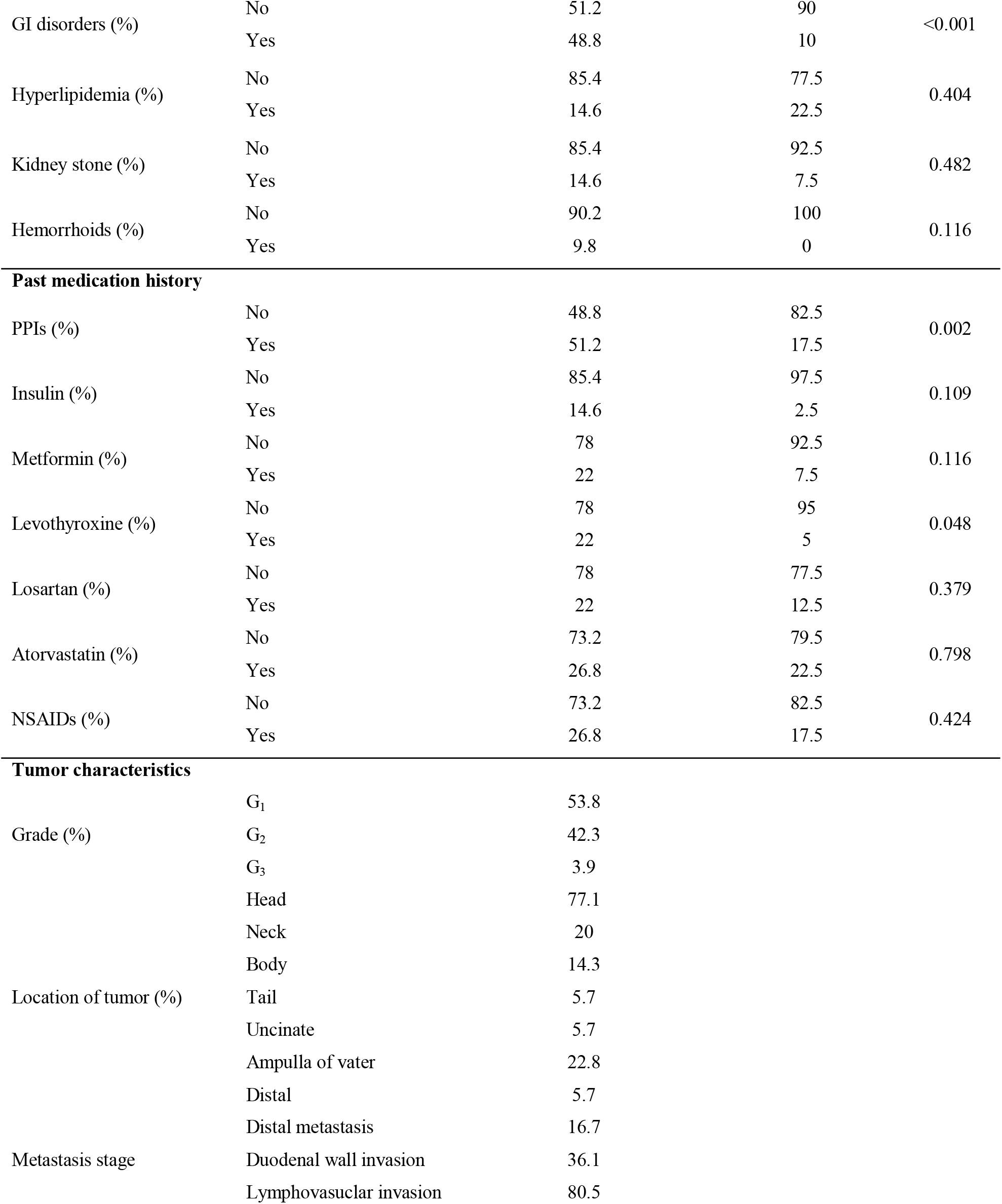
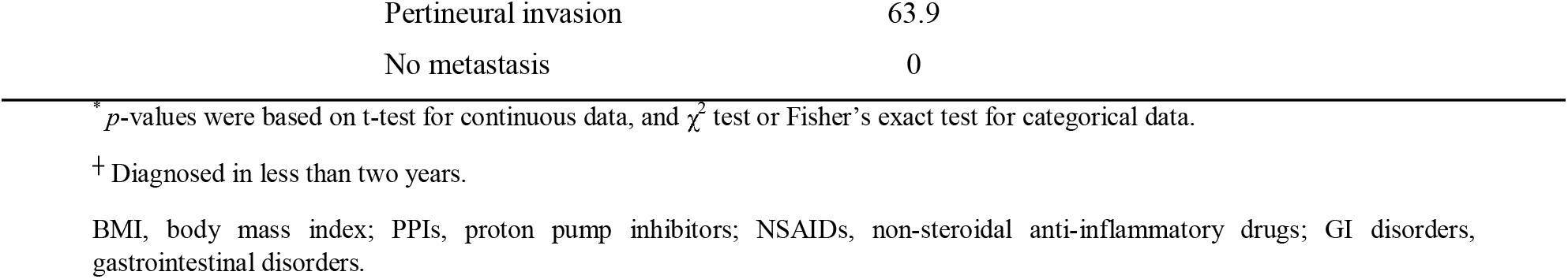
Distribution of demographic parameters among pancreatic cancer cases and non-cancer controls.

### Salivary and blood markers differences among groups

*P. gingivalis* was more prevalent in saliva from patients than controls. It was detected in approximately 61% (25 out of 41) of the patient group but was found only in 22.5% (nine out of 43) of the controls. Furthermore, a significant difference was observed in the number of *P. gingivalis* cells in salivary samples between patients and controls (*p*=0.034). Similarly, *A. actinomycetemcomitans* was more prevalent among patients, although no significant difference was found between the two groups (*p*=0.128). Additionally, miR-155 were overexpressed, with a significant increase (*p*=0.034) in copy numbers observed in patients with PC compared to non-cancer individuals. Conversely, despite the overexpression of miR-21 in PC group, no statistically significant distinctions found between the two groups (*p*=0.175) (Table 2). Following the adjustment for potential confounders, a moderate and statistically significant association emerged between the disease status of subjects and the composition of microbial loads, as well as the copy number of miRNAs (R2=0.035, *p*=0.046) (Figure 1).

**Table 2:**
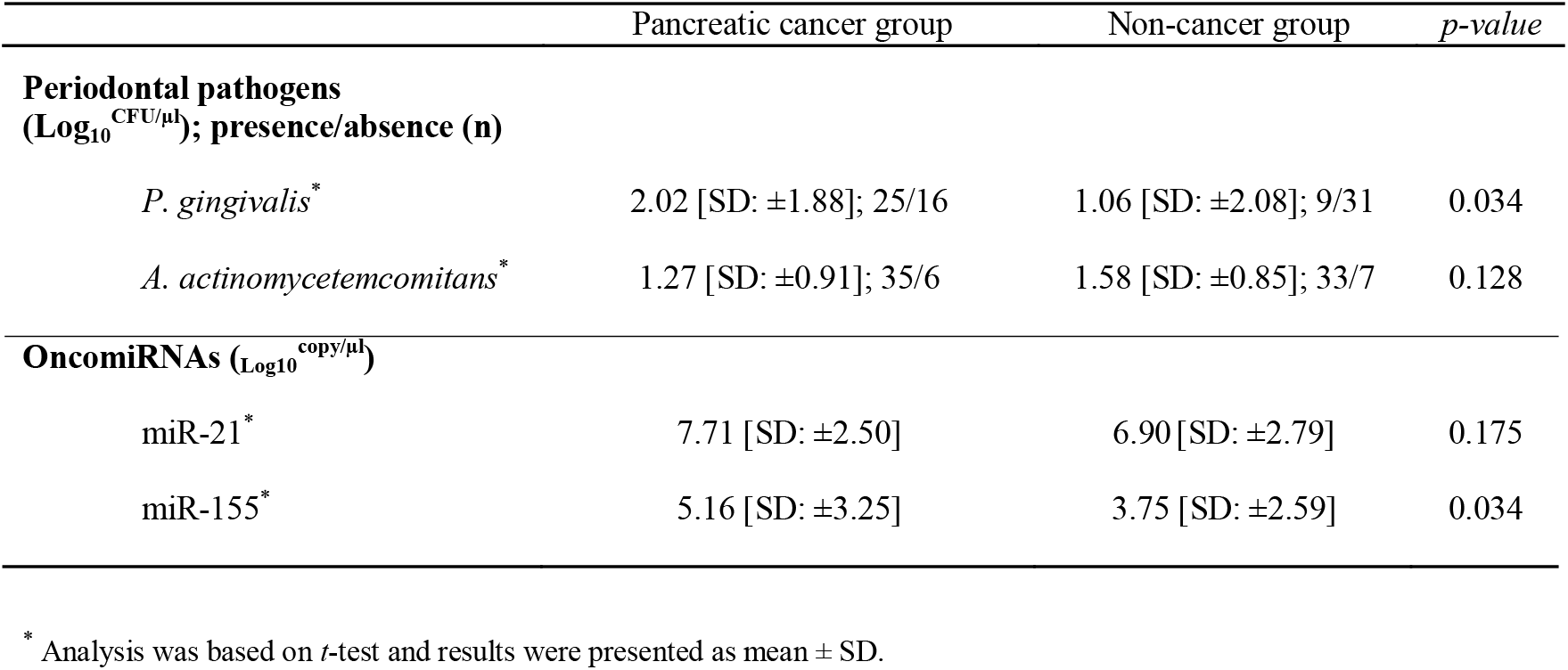
Absolute value of periodontal pathogens in saliva and oncomiRNAs in blood samples of PC patients and controls assessed by real-time PCR.

| Pancreatic cancer group | Non-cancer group | $p$-value |
| --- | --- | --- |

|  |  |  |  |
| --- | --- | --- | --- |
| **Periodontal pathogens**<br>(Log<sub>10</sub><sup>CFU/μl</sup>); presence/absence (n) |  |  |  |
| *P. gingivalis* * | 2.02 [SD: ±1.88]; 25/16 | 1.06 [SD: ±2.08]; 9/31 | 0.034 |
| *A. actinomycetemcomitans* * | 1.27 [SD: ±0.91]; 35/6 | 1.58 [SD: ±0.85]; 33/7 | 0.128 |
| **OncomiRNAs (Log<sub>10</sub><sup>copy/μl</sup>)** |  |  |  |
| miR-21* | 7.71 [SD: ±2.50] | 6.90 [SD: ±2.79] | 0.175 |
| miR-155* | 5.16 [SD: ±3.25] | 3.75 [SD: ±2.59] | 0.034 |

\* Analysis was based on *t*-test and results were presented as mean ± SD.

**Figure1.**
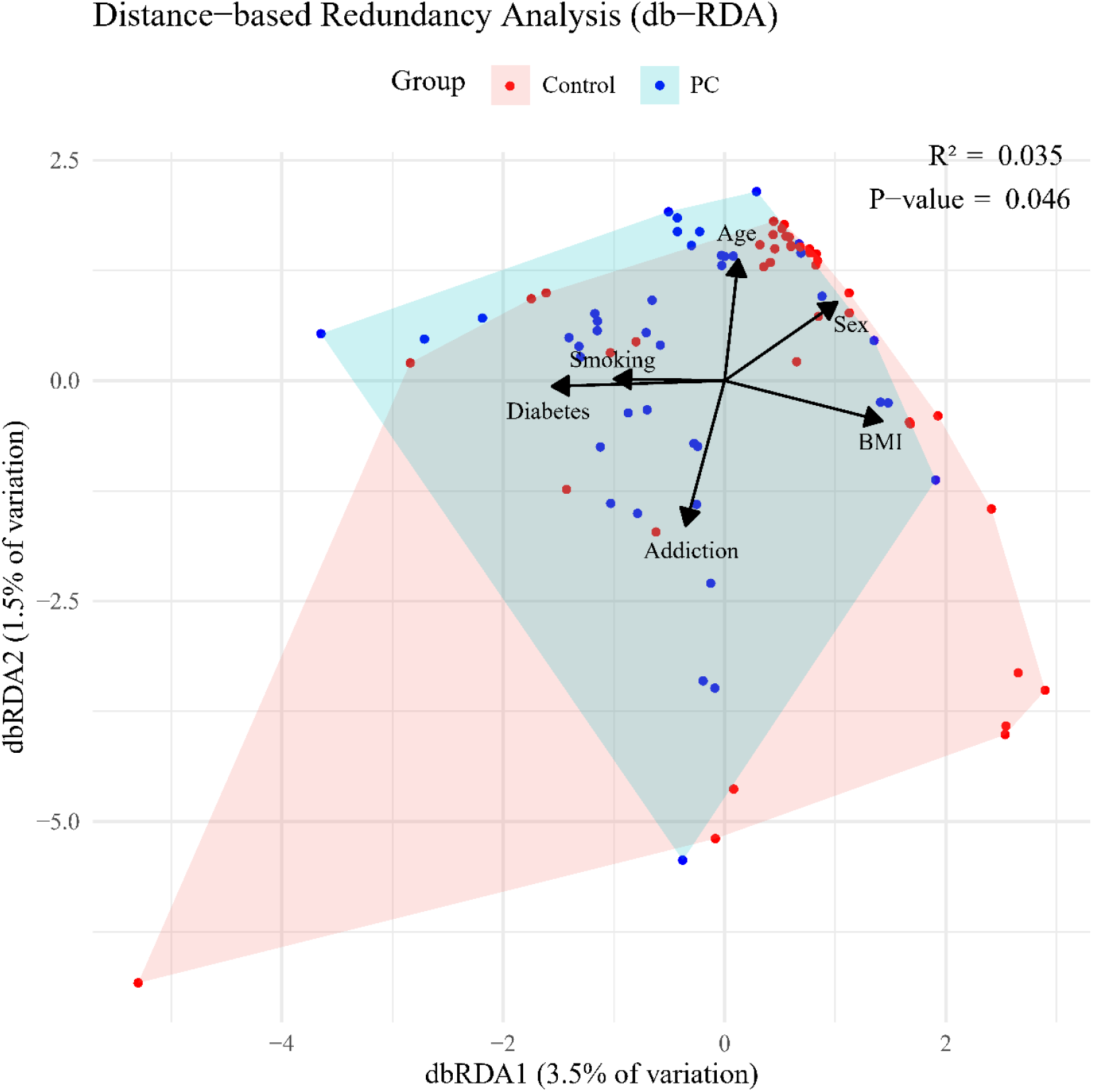
Bray-Curtis distance-based redundancy analysis (dbRDA) based on salivary loads of *Porphyromonas gingivalis* and *Aggregatibacter actinomycetemcomitans*, along with circulating miR-21 and miR-155 copy number variation, in pancreatic cancer (PC) versus control groups.

### Biomarker associations, stratified risk analyses, and predictive model performance

Elevated levels of *P. gingivalis* were associated with a higher risk of PC (adjusted OR=1.43 and 95% CI 1.04 to 1.96). Overexpression of miR-21 was also associated with an increased risk of PC (adjusted OR=1.33 and 95% CI 1.04 to 1.70). In contrast, loads of *A. actinomycetemcomitans* and miR-155 were not significantly associated with the risk of PC (Table S2). In analyses stratified by age, adjusted OR for *P. gingivalis* and miR-155 in relation to PC risk were 1.67 (95% CI 1.17 to 2.39) and 1.44 (95% CI 1.13 to 1.84) among individuals younger than years old, respectively. Among females, enrichment of *P. gingivalis* was associated with a higher risk of PC (adjusted OR=2.31 and 95% CI 0.98 to 5.47) compared to males (adjusted OR=1.36 and 95% CI 0.98 to 1.90). In analyses stratified by smoking status, no significant differences in the risk of PC were observed among ever smokers for the selected biomarkers. Notably, among individuals with type 2 diabetes, greater abundance of *A. actinomycetemcomitans* was associated with an approximately 3-fold increased risk of PC (adjusted OR = 3.66, 95% CI 0.47 to 6.68), compared to the risks associated with elevated levels of *P. gingivalis* (adjusted OR = 1.16, 95% CI 0.60 to 2.24) (Table 3). However, within non-diabetic individuals, only *P. gingivalis* related to the rise of PC risk (adjusted OR = 1.67, 95% CI 1.05 to 2.66).

**Table 3:**
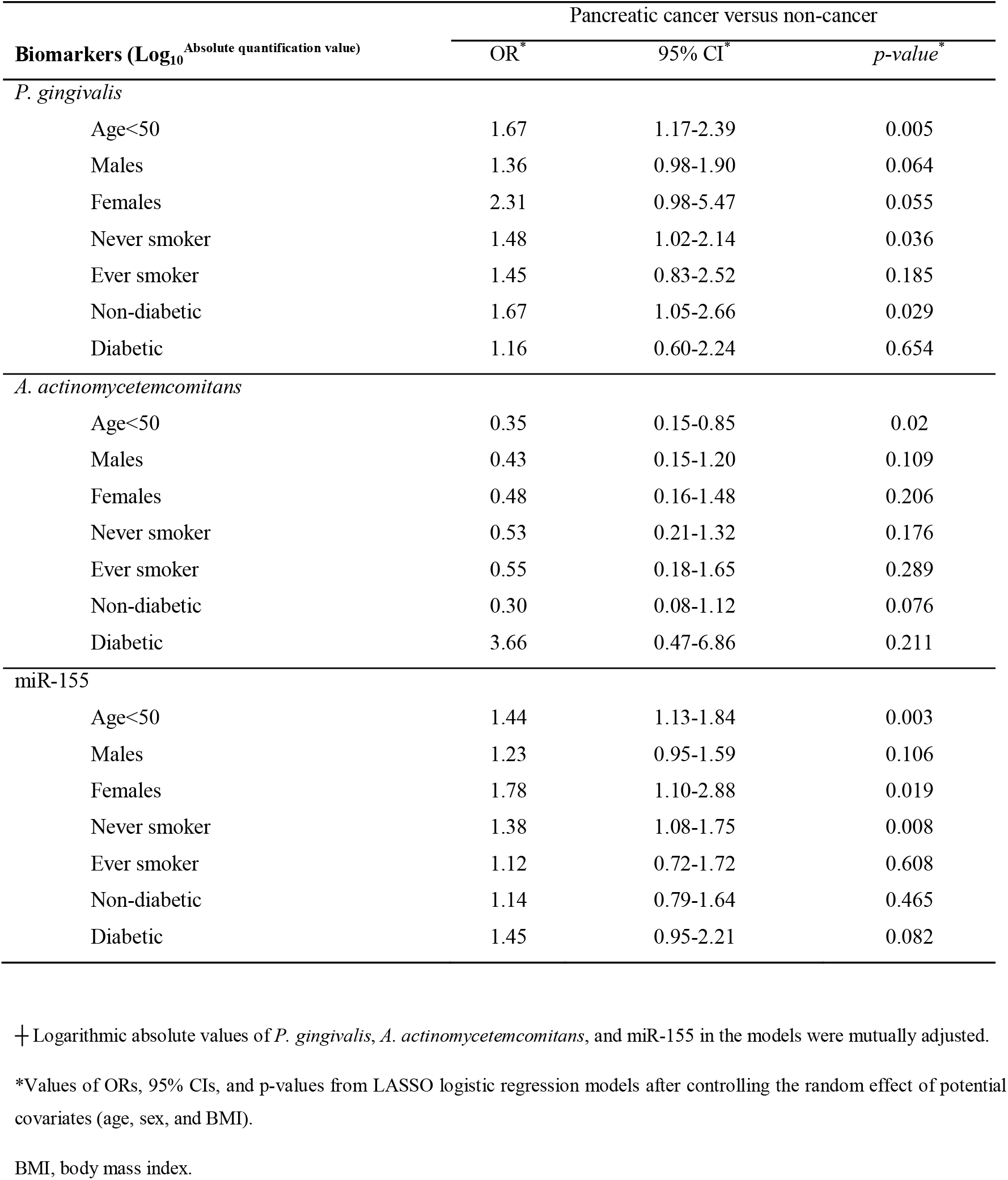
Selected biomarkers^┼^ and risk of pancreatic cancer, stratified by age, sex, smoking, and diabetes between pancreatic cancer cases and non-cancer controls.

To evaluate the performance of the biomarkers of interest, five models were constructed (Figure 2). The first model, based on LASSO feature selection, achieved an AUC of 0.080 (95% CI 0.707 to 0.909), with a sensitivity of 73.1% and a specificity of 86.8%. Intriguingly, model 4, which incorporated our four main biomarkers along with diabetes status, demonstrated the highest AUC of 0.878 (95% CI 0.802 to 0.955), indicating that these factors together predict the occurrence of PC with the highest accuracy (0.835). Meanwhile, model 5 exhibited the highest sensitivity (90.2%), and model 2 showed the highest specificity (92.1%) (Figure 2 and Table S3).

**Figure 2.**
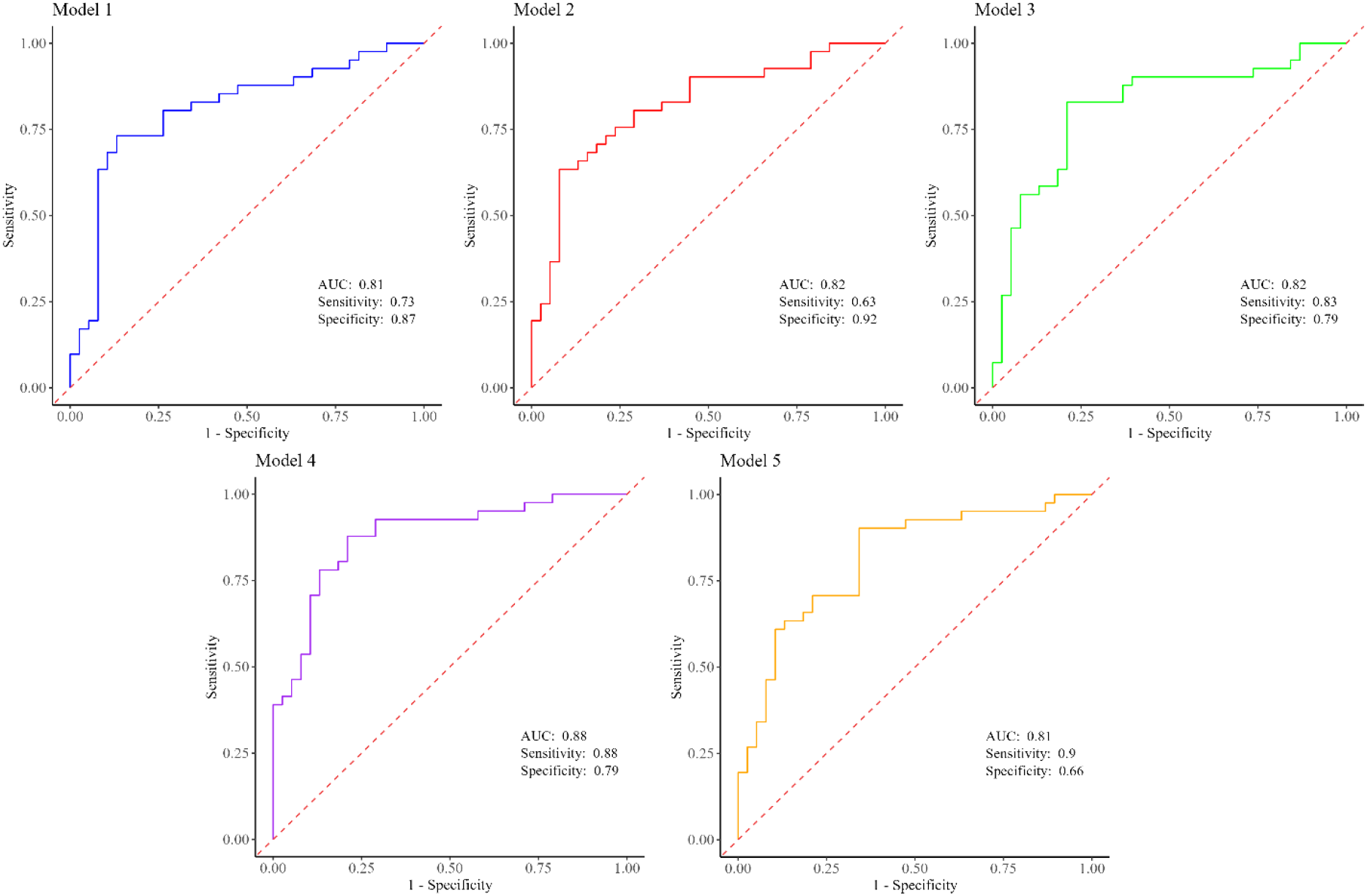
Predictive performance estimates between PC patients and non-cancer controls for model 1 (Pg+Aa+miR-155+BMI), model 2 (Pg+Aa+miR-21+miR-155+BMI), model 3 (Pg+Aa+miR-21+miR-155), model 4 (Pg+Aa+miR-21+miR-155+diabetes), and model 5 (Pg+Aa+miR-21+miR-155+smoking). PC, pancreatic cancer; Pg, *Porphyromonas gingivalis*; Aa, *Aggregatibacter actinomycetemcomitans*; BMI, body mass index.

## Discussion

*P. gingivalis* and *A. actinomycetemcomitans* play pivotal roles as keystone pathogens within the oral microbiome, contributing significantly to the onset of periodontal disease and dental deterioration (25). Prior research has consistently highlighted that a history of periodontal disease and tooth loss correlates with a heightened likelihood of developing PC (13-15, 26, 27). The present case-control investigation revealed a significant association between the salivary load of *P. gingivalis* and the expression level of miR-155 in individuals suffering from PC compared to controls. According to our results, among the two examined pathogens, *P. gingivalis* was significantly associated with a notable OR of 1.43 after adjusting for potential confounders including age, gender, diabetes status, and smoking. Interestingly, higher levels of *P. gingivalis* in females were related to the highest increase in the risk of PC by nearly two-fold. Subsequently, individuals younger than 50 years old and non-diabetic individuals were at a heightened risk compared to other subgroups following an enrichment of *P. gingivalis*. Among studies conducted on the oral microbiome, three nested case-control studies have investigated the relationship between PC and *P. gingivalis* (13, 18, 19). Two of these studies focused on samples obtained from the mouthwash of patients before the onset of PC (18, 19). According to previous findings, the risk of developing PC in the presence of *P. gingivalis* increased by 1.60 (18). Additionally, never smokers and never drinkers were reported to have the highest risk of developing the disease for carriage of *P. gingivalis* (18, 19). In another prospective study conducted on blood samples from 405 individuals prior to developing PC, it was revealed that in samples where the level of antibodies against *P. gingivalis* exceeded 200 ng/ml, the risk of PC doubled (13). Our findings are consistent with prior research indicating an association between the presence of *P. gingivalis* and the incidence of PC. However, in previous studies, the odds ratio was either based on the presence versus absence or relative abundance of this pathogen, whereas in our studies, the observed association was based on the number of *P. gingivalis* cells present in salivary samples.

Another keystone pathogen in the oral cavity is *A. actinomycetemcomitans*, which has yielded conflicting results. While one study revealed an association with an OR of 2.20 (18), both our investigation and other research failed to show a significant correlation with the risk of developing PC (13, 19). Previous investigations have explored the intriguing aspect of assessing the presence of this periodontal pathogen in relation to PC risk within different subgroups. Noteworthy odds ratios were only significant within the subgroup of individuals who reported consuming alcohol, where the presence of bacteria was linked to an almost threefold increase in PC risk (18). Notably, in our study, the majority of the population did not report a history of alcohol consumption according to self-report questionnaires. In our subgroup analysis, individuals with diabetes were discovered to have an increased susceptibility to PC associated with the sheer number of *A. actinomycetemcomitans* cells in the oral cavity (OR=3.66). The comparatively low prevalence of *A. actinomycetemcomitans* in oral samples posed a notable challenge, as highlighted in previous studies. However, in our study, we addressed this issue effectively by employing absolute quantification techniques.

In the past few decades, there has been a strenuous effort in scientific research focusing on the oral microbiome to pinpoint high-sensitivity and high-specificity biomarkers for the early and non-invasive detection of PC (28-32). Concurrently, research endeavors have delved into exploring circulating miRNAs for similar diagnostic objectives (33, 34). To the best of our knowledge, for the first time, we combined the absolute abundance of two periodontal pathogens with copy number of two oncomiRNAs to develop predictive models for the detection of PC. Diagnosis of PC at an early stage remains a significant challenge in mitigating the impact of this malignancy. Currently, the only Food and Drug Administration (FDA)-approved biomarker for PC is serum CA19-9, primarily utilized for monitoring disease progression rather than screening, due to its insufficient sensitivity and specificity (35). Therefore, developing non-invasive markers for either screening or early detection of the disease has been catapulting interest recently. In our study, we built a predictive model that accurately predicts PC solely based on a characteristic panel of four main biomarkers. Moreover, the diagnostic power of *P. gingivalis* and *A. actinomycetemcomitans* has not been reported in any previous study. Nevertheless, to date, numerous studies have been conducted on the diagnostic power of oral microbiome profiles in PC. In a study by Farrell and colleagues, which included 28 individuals with PC and 28 healthy controls (HCs), an AUC of 0.90 with a 96.4% sensitivity and 82.1% specificity was reported for *Neisseria elongata* and *Streptococcus mitis* (28). In another model developed using data from 20 individuals with PC and 16 HCs, the ten genera identified together yielded an AUC of 0.916 (29). The third study on 30 PC patients and 25 HCs, using a combination of four genera (*Porphyromonas, Fusobacterium, Haemophilus*, and *Leptotrichia*), achieved an AUC of 0.802 with a 77.1% sensitivity and 78.6% specificity (30). In a study by Nagata and colleagues on 47 PC patients and 235 controls, two sets of oral microbiome combinations (mostly composed of *Streptococcus* species) were identified with AUC of 0.80 and 0.82 (31). Many studies have also investigated the diagnostic power of circulating miR-21 and miR-155 for the detection of PC, yet each miRNA individually yielded an AUC not surpassing 0.78, showcasing low sensitivity but notable specificity (33, 34). Likewise, upon integrating miR-21 into our LASSO-based model (model 1), the specificity escalated to 92.1%, albeit at the cost of reduced sensitivity to 63.4%. Intriguingly, within our models, the combination of four primary biomarkers along with diabetes status, demonstrated the most elevated AUC of 0.878, exhibiting a specificity of 78.9% and a sensitivity of 87.8%.

Despite implementing an innovative methodology for biomarker discovery, our study was constrained by several limitations. The primary limitation of the current study was the one-time sample collection, which restricted our capacity to capture potential temporal dynamics within the relationship among the main variables and PC. Another limitation of this study was the small sample size, which precluded the validation and robustness evaluation of our predictive models. The final limitation was that we did not have any data on history of periodontal diseases, preventing us from adjusting for it as a potential confounding factor.

## Conclusions

In summary, for the first time, this study represented the exploration of the collective diagnostic effectiveness of *P. gingivalis, A. actinomyctemcomitans*, miR-21, and miR-155, demonstrating heightened disease specificity and accuracy. The results implied that the proposed panel of biomarkers shows the potential to enhance non-invasive PC detection, supplementing existing markers and paving the way for cost-effective PC screening and monitoring. Furthermore, we believe that these identified biomarkers might extend beyond diagnostic purposes, offering potential for varied applications in prevention and therapeutic strategies.

## Supplementary Materials

Table S1: Primer sequences for a quantitative analysis by qPCR of periodontal pathogens and oncomiRNAs, Table S2: Multivariate analysis of risk assessment for prominent biomarkers between individuals with pancreatic cancer and non-cancer controls, Table S3: Performance estimates of the constructed models based on salivary and blood biomarkers.

## Author contributions

Writing–original draft, Z.H.; Visualization Z.H., M.O., and H.E.; Investigation, Z.H., S.M., and H.E.; Formal analysis, Z.H., M.O., S.M., and H.E.; Data curation, Z.H., M.O., and H.H.; Methodology, Z.H., A.S., M.O., M.H., and H.H.; Writing–review & editing, A.S., S.M., H.E., M.H., and H.H.; Validation, V.P., M.H., and H.H.; Resources, V.P., S.M., H.E., M.H., and H.H.; Project administration, M.H., V.P., and H.H.; Software, M.O.; Supervision, M.H. and V.P.; Funding acquisition, V.P.; Conceptualization, H.H.

## Institutional Review Board Statement

This study was reviewed and approved by the Institutional Ethical Review Committee of the School of Medicine at Shahid Beheshti University of Medical Sciences (No.IR.SBMU.MSP.REC.1402.184).

## Informed Consent Statement

Informed consent was obtained from all subjects involved in the study.

## Data Availability Statement

All data generated or analyzed during this study are included in this article.

## Acknowledgments

The authors would like to express their sincere gratitude to the Endoscopy Department at Taleghani Hospital for their invaluable cooperation and support during this study. They extend their heartfelt appreciation to the esteemed members of the Research Institute for Gastroenterology and Liver Diseases at Shahid Beheshti University of Medical Sciences in Tehran, Iran. Additionally, the authors wish to acknowledge Professor Mohammad Reza Zali for his unwavering support throughout the project. They also extend their sincere thanks to Professor Masoumeh Douraghi from the School of Public Health at Tehran University of Medical Sciences, along with the dedicated team at the Dental School of Shahid Beheshti University of Medical Sciences, for providing standard strain of periodontal pathogens.

## References

1. Ferlay J, Colombet M, Soerjomataram I, Parkin DM, Piñeros M, Znaor A, Bray F. Cancer statistics for the year 2020: An overview. Int J Cancer. 2021.

2. Hesami Z, Olfatifar M, Sadeghi A, Zali MR, Mohammadi-Yeganeh S, Habibi MA, et al. Global Trend in Pancreatic Cancer Prevalence Rates Through 2040: An Illness-Death Modeling Study. Cancer Med. 2024;13(20):e70318.

3. Hezel AF, Kimmelman AC, Stanger BZ, Bardeesy N, Depinho RA. Genetics and biology of pancreatic ductal adenocarcinoma. Genes Dev. 2006;20(10):1218–49.

4. Li D, Xie K, Wolff R, Abbruzzese JL. Pancreatic cancer. Lancet. 2004;363(9414):1049–57.

5. Whitcomb DC. Inflammation and Cancer V. Chronic pancreatitis and pancreatic cancer. Am J Physiol Gastrointest Liver Physiol. 2004;287(2):G315–9.

6. Jemal A, Siegel R, Ward E, Hao Y, Xu J, Murray T, Thun MJ. Cancer statistics, 2008. CA Cancer J Clin. 2008;58(2):71–96.

7. Ries L, Melbert D, Krapcho M, Stinchcomb D, Howlader N, Horner M. SEER Cancer Statistics Review, 1975–2005, National Cancer Institute. Bethesda, MD, SEER Cancer Statistics Review, 1975–2005, National Cancer Institute. Bethesda, MD. 2007.

8. Rawla P, Sunkara T, Gaduputi V. Epidemiology of Pancreatic Cancer: Global Trends, Etiology and Risk Factors. World J Oncol. 2019;10(1):10–27.

9. Wood LD, Yurgelun MB, Goggins MG. Genetics of Familial and Sporadic Pancreatic Cancer. Gastroenterology. 2019;156(7):2041–55.

10. Ohara Y, Valenzuela P, Hussain SP. The interactive role of inflammatory mediators and metabolic reprogramming in pancreatic cancer. Trends Cancer. 2022;8(7):556–69.

11. Aas JA, Paster BJ, Stokes LN, Olsen I, Dewhirst FE. Defining the normal bacterial flora of the oral cavity. J Clin Microbiol. 2005;43(11):5721–32.

12. Zhang Y, Niu Q, Fan W, Huang F, He H. Oral microbiota and gastrointestinal cancer. Onco Targets Ther. 2019;12:4721–8.

13. Michaud DS, Joshipura K, Giovannucci E, Fuchs CS. A prospective study of periodontal disease and pancreatic cancer in US male health professionals. J Natl Cancer Inst. 2007;99(2):171–5.

14. Hujoel PP, Drangsholt M, Spiekerman C, Weiss NS. An exploration of the periodontitis-cancer association. Ann Epidemiol. 2003;13(5):312–6.

15. Stolzenberg-Solomon RZ, Dodd KW, Blaser MJ, Virtamo J, Taylor PR, Albanes D. Tooth loss, pancreatic cancer, and Helicobacter pylori. Am J Clin Nutr. 2003;78(1):176–81.

16. Alkharaan H, Lu L, Gabarrini G, Halimi A, Ateeb Z, Sobkowiak MJ, et al. Circulating and Salivary Antibodies to Fusobacterium nucleatum Are Associated With Cystic Pancreatic Neoplasm Malignancy. Front Immunol. 2020;11:2003.

17. Yu J, Ploner A, Chen MS, Zhang J, Sandborgh-Englund G, Ye W. Poor dental health and risk of pancreatic cancer: a nationwide registry-based cohort study in Sweden, 2009-2016. Br J Cancer. 2022;127(12):2133–40.

18. Fan X, Alekseyenko AV, Wu J, Peters BA, Jacobs EJ, Gapstur SM, et al. Human oral microbiome and prospective risk for pancreatic cancer: a population-based nested case-control study. Gut. 2018;67(1):120–7.

19. Petrick JL, Wilkinson JE, Michaud DS, Cai Q, Gerlovin H, Signorello LB, et al. The oral microbiome in relation to pancreatic cancer risk in African Americans. Br J Cancer. 2022;126(2):287–96.

20. Chang TC, Mendell JT. microRNAs in vertebrate physiology and human disease. Annu Rev Genomics Hum Genet. 2007;8:215–39.

21. Hwang HW, Mendell JT. MicroRNAs in cell proliferation, cell death, and tumorigenesis. Br J Cancer. 2006;94(6):776–80.

22. Seyhan AA. Circulating microRNAs as Potential Biomarkers in Pancreatic Cancer-Advances and Challenges. Int J Mol Sci. 2023;24(17).

23. Metcalf GAD. MicroRNAs: circulating biomarkers for the early detection of imperceptible cancers via biosensor and machine-learning advances. Oncogene. 2024;43(28):2135–42.

24. Mohammadi-Yeganeh S, Paryan M, Mirab Samiee S, Soleimani M, Arefian E, Azadmanesh K, et al. Development of a robust, low cost stem-loop real-time quantification PCR technique for miRNA expression analysis. Molecular Biology Reports. 2013;40(5):3665–74.

25. Costalonga M, Herzberg MC. The oral microbiome and the immunobiology of periodontal disease and caries. Immunol Lett. 2014;162(2 Pt A):22–38.

26. Ahn J, Segers S, Hayes RB. Periodontal disease, Porphyromonas gingivalis serum antibody levels and orodigestive cancer mortality. Carcinogenesis. 2012;33(5):1055–8.

27. Hiraki A, Matsuo K, Suzuki T, Kawase T, Tajima K. Teeth loss and risk of cancer at 14 common sites in Japanese. Cancer Epidemiol Biomarkers Prev. 2008;17(5):1222–7.

28. Farrell JJ, Zhang L, Zhou H, Chia D, Elashoff D, Akin D, et al. Variations of oral microbiota are associated with pancreatic diseases including pancreatic cancer. Gut. 2012;61(4):582–8.

29. Chen T, Li X, Li G, Liu Y, Huang X, Ma W, et al. Alterations of commensal microbiota are associated with pancreatic cancer. Int J Biol Markers. 2023;38(2):89–98.

30. Lu H, Ren Z, Li A, Li J, Xu S, Zhang H, et al. Tongue coating microbiome data distinguish patients with pancreatic head cancer from healthy controls. J Oral Microbiol. 2019;11(1):1563409.

31. Nagata N, Nishijima S, Kojima Y, Hisada Y, Imbe K, Miyoshi-Akiyama T, et al. Metagenomic Identification of Microbial Signatures Predicting Pancreatic Cancer From a Multinational Study. Gastroenterology. 2022;163(1):222–38.

32. Kartal E, Schmidt TSB, Molina-Montes E, Rodríguez-Perales S, Wirbel J, Maistrenko OM, et al. A faecal microbiota signature with high specificity for pancreatic cancer. Gut. 2022;71(7):1359–72.

33. Wang J, Chen J, Chang P, LeBlanc A, Li D, Abbruzzesse JL, et al. MicroRNAs in plasma of pancreatic ductal adenocarcinoma patients as novel blood-based biomarkers of disease. Cancer Prev Res (Phila). 2009;2(9):807–13.

34. Qu K, Zhang X, Lin T, Liu T, Wang Z, Liu S, et al. Circulating miRNA-21-5p as a diagnostic biomarker for pancreatic cancer: evidence from comprehensive miRNA expression profiling analysis and clinical validation. Scientific reports. 2017;7(1):1692.

35. Goonetilleke KS, Siriwardena AK. Systematic review of carbohydrate antigen (CA 19-9) as a biochemical marker in the diagnosis of pancreatic cancer. Eur J Surg Oncol. 2007;33(3):266–70.

