## Supplementary material for "Interplay of *Porphyromonas gingivalis* and *Aggregatibacter actinomycetemcomitans* along with Circulating miR-21 and miR-155 as Potential Biomarkers for Pancreatic Cancer": Supplamantry File 1

Table S1: Primer sequences for a quantitative analysis by qPCR of periodontal pathogens and oncomiRNAs

| | Oligonucleotide 5′ - 3′ | Product size | Reference |
|---|---|---|---|
| **Periodontal pathogens** | | 16SrRNA | |
| *P. gingivalis* | F: 5′- TGTAGATGACTGATGGTGAAAACC -3′<br>R:5′- ACGTCATCCACACCTTCCTC -3′ | 198 bp | (1) |
| *A. actinomycetemcomitans* | F: 5′- ATTGGGGTTTAGCCCTGGT -3′<br>R:5′- GGCACAAACCCATCTCTGA-3′ | 250 bp | (2) |
| **OncomiRNAs** | | | |
| miR-21 | F: 5′- GTATACTAGCTTATCAGACTG -3′<br>R:5′- GTGCAGGGTCCGAGGT -3′ | 79 bp | (3) |
| miR-155 | F: 5′- GTATACTTAATGCTAATTGTG -3′<br>R:5′- GTGCAGGGTCCGAGGT -3′ | 80 bp | (4) |

Table S2: Multivariate analysis of risk assessment for prominent biomarkers┼ between individuals with pancreatic cancer and non-cancer controls

| | Pancreatic cancer verses non-cancer | | |
|---|---|---|---|
| **Biomarkers** | OR* | 95% CI* | *p* value* |
| *P. gingivalis* | | | |
| $Log_{10}^{Absolute\ quantification\ value}$ | 1.43 | 1.04-1.96 | 0.024 |
| *A. actinomycetemcomitans* | | | |
| $Log_{10}^{Absolute\ quantification\ value}$ | 0.68 | 0.34-1.36 | 0.282 |
| miR-21 | | | |
| $Log_{10}^{Absolute\ quantification\ value}$ | 1.33 | 1.04-1.70 | 0.019 |
| miR-155 | | | |
| $Log_{10}^{Absolute\ quantification\ value}$ | 1.15 | 0.92-1.44 | 0.190 |

CI, confidence interval; OR, odds ratio.

┼Logarithmic absolute values of *P. gingivalis*, *A. actinomycetemcomitans*, miR-21 and miR-155 in the models were mutually adjusted.

*Values of ORs, 95% CIs and p values from logistic regression models after controlling the random effect of potential covariates (age, sex, smoking status, and diabetes).

Table S3: Performance estimates of the constructed models based on salivary and blood biomarkers

| Models | Features | AUC (95% CI) | Accuracy (%) | Specificity (%) | Sensitivity (%) |
|---|---|---|---|---|---|
| 1 | Pg+Aa+miR-155+BMI | 0.808 (0.707-0.909) | 79.7 | 86.8 | 73.1 |
| 2 | Pg+Aa+miR-21+miR-155+BMI | 0.817 (0.722-0.912) | 77.2 | 92.1 | 63.4 |
| 3 | Pg+Aa+miR-21+miR-155 | 0.815 (0.716-0.913) | 81.0 | 78.9 | 82.9 |
| 4 | Pg+Aa+miR-21+miR-155+Diabetes | 0.878 (0.802-0.955) | 83.5 | 78.9 | 87.8 |
| 5 | Pg+Aa+miR-21+miR-155+Smoking | 0.814 (0.719-0.91) | 78.4 | 65.7 | 90.2 |

CI, confidence interval; AUC, area under the curve; Pg, *Porphyromonas gingivalis*; Aa, *Aggregatibacter actinomycetemcomitans*.

### References


1. Hu Z, Zhang Y, Li Z, Yu Y, Kang W, Han Y, et al. Effect of Helicobacter pylori infection on chronic periodontitis by the change of microecology and inflammation. Oncotarget. 2016;7(41):66700-12.
2. Rudney JD, Chen R, Pan Y. Endpoint quantitative PCR assays for Bacteroides forsythus, Porphyromonas gingivalis, and Actinobacillus actinomycetemcomitans. J Periodontal Res. 2003;38(5):465-70.
3. Houri H, Ghalavand Z, Faghihloo E, Fallah F, Mohammadi-Yeganeh S. Exploiting yoeB-yefM toxin-antitoxin system of Streptococcus pneumoniae on the selective killing of miR-21 overexpressing breast cancer cell line (MCF-7). Journal of Cellular Physiology. 2020 Mar;235(3):2925-36.
4. Kia V, Paryan M, Mortazavi Y, Biglari A, Mohammadi-Yeganeh S. Evaluation of exosomal miR-9 and miR-155 targeting PTEN and DUSP14 in highly metastatic breast cancer and their effect on low metastatic cells. Journal of cellular biochemistry. 2019 Apr;120(4):5666-76.
